# Ovarian cysts and granulosa cell tumors develop after sublethal total body irradiation in mice

**DOI:** 10.1101/2021.07.08.450892

**Authors:** Shiyun Xiao, Yao Xiao, Xiaohong Xu, Wen Zhang, Mandi M. Murph, Nancy R. Manley

## Abstract

**Background:** Localized and total body irradiation are used to treat certain cancers and also used prior to transplantation of stem cells or organs. However, the use of radiation also induces collateral damage to the cells of healthy tissue. Although the acute damage of radiation to oocytes is well known, the long-term effects induced by radiation to stromal cells and their relationship with age are still unclear.

**Methods:** A total of 206 two-month-old female mice were whole-body exposed to gamma rays at doses of 0, 0.5, 1, 2, or 4 Gy, respectively. The mice were sacrificed at 3.5, 9, 12, or 18 months of age and pathological changes including cysts and tumors were assessed in the ovary and other organs.

**Results:** The overall incidence of visible pathological changes of mice receiving irradiation was 33.7% in the ovary, but much lower in the liver, spleen, lung, thymus, and skin. Among these, the ovarian cyst formation rate was 24.7%, and tumor lesions were 10.2%, respectively, compared to 5% cyst formation and no tumor lesions among control, unirradiated mice. Statistical analysis showed that cyst formation was age, but not dose-dependent, whereas the formation of tumor lesions was dependent on both age and radiation dose. Pathology analysis indicated that most ovarian cysts originated from follicles and both tumor lesions analyzed originated from granulosa cells.

**Conclusion:** Ovaries are highly susceptible to the effects of radiation. Long-term damage is increased after total body irradiation in mice, manifested by higher incidences of cyst formation and tumor lesions. The ovarian stromal-derived granulosa cells might play an essential role in these changes.

## Introduction

Ionizing irradiation is formed by charged energy particles, which can directly or indirectly cause damage to the DNA when it passes through the body [1]. Following radiation exposure, DNA repair mechanisms are initiated in the cell. In general, a lower-absorbed dose of radiation causes repairable DNA damage, whereas a higher-absorbed dose causes significant DNA damage and tends to kill the irradiated cells [2, 3]. Proper DNA repair restores the cells and tissues to normal; mis-repaired or un-repaired DNA causes cell death, which can lead to damage or failure of the tissue or organ. In some cases, the cells with the mis-repaired or un-repaired DNA can survive after irradiation, and can be a substrate for tumor/cancer development [2, 4].

Different organs, tissues, and cells, and even the same type of cell in a different phase, appear to have different radiation sensitivity [5-8]. The ovary is the female gonad and the primary reproductive organ, which is composed of an outer cortex and an inner medulla, and the port of entry into the ovary is known as the ovarian hilus [9]. The outer cortex contains developing follicles at various stages and is the most radiation-sensitive part [10]. The cortical primordial follicle pool is nearly destroyed in young adult female mice two weeks after a single total body exposure to the dose of 0.1 Gy [11]. The complete destruction of the primordial follicle pool was reported in neonatal mice five days after exposure to doses as low as 0.45 Gy [12]. In contrast, the partial primary, secondary follicles, and large follicles still survived, showing more resistance to the radiation than the primordial follicles [7, 10, 12, 13]. The inner medulla contains the major blood and lymphatic vessels connected to the hilus and cortex stromal compartment, known as the ovarian niche, which supports the development of oocyte follicles. These are largely considered to be radiation resistant. However, the radiation damage to the follicles may destroy not only the cortical construction but also the medulla stromal cells. It is these changes that usually cause ovarian failure or dysfunction and therefore lead to premature reproductive senescence or permanent menopause after irradiation [14-16].

Ovarian cysts are a frequent problem among reproductive-aged females undergoing ovulation cycles and aging rodents with hormone variations [17, 18]. During each cycle, mature follicles forms lumps on the ovary and rupture at ovulation, releasing an oocyte. Rodents have an estrous cycle that lasts 4 to 5 days with mice ovulating an average of 11 oocytes. Fluid-filled cysts stem from mature follicles that do not rupture to release the oocyte after the period of follicular growth and development [19].

Granulosa cells play a critical role in this dynamic process. In order for successful ovulation and follicular development to occur, granulosa cells surrounding oocytes must replicate to form multiple layers and continually alter their morphology. In fact, a major characteristics of mature follicle formation includes a fluid-filled vesicle that materializes as a result of proper granulosa cells’ organization [19]. If the mature follicle does not rupture appropriately, a fluid-filled cyst remains, however, cysts are considered physiological changes and often resolve naturally over 1-3 months [20, 21]. Interestingly, granulosa cells secrete growth-promoting factors along with their receptors. As such, they are the main source of estradiol, insulin-like growth factor and others that stimulate follicular growth and survival [22].

Exposure to radiation is a known cause of cancer, whereas reducing the overall number of lifetime ovulations, which occurs with pregnancy or the use of oral contraception, decreases the risk of ovarian cancer [23, 24]. Most previous studies on ovarian damage induced by radiation have focused on the acute effects of follicle deletion and its related mechanisms, few have focused on niche changes and long-term radiation effects on the ovary. Thus, our study provided an opportunity to investigate the ovary for relationships between radiation damage, cyst formation and tumor development.

In this study, two-month-old female mice were exposed to gamma rays (Co-60) at sub-lethal doses ranging from 0.5 to 4 Gy, and radiation effects were analyzed at 3.5, 9, 12, and 18 months of age by comparison to 0 Gy control animals. We previously reported late effects of radiation on the immune system [25]; in the course of that study, we also observed pathological changes like cysts and tumors increasing significantly in the mouse ovary, but not in other organs. Our results herein demonstrate that the ovary is a highly radiation-sensitive organ in mice, and damage sustained after a single total body irradiation (TBI) increases ovarian cysts and tumor lesions. Visible ovarian cysts (the size > 1 mm^3^) were observed starting at 9 months of age (7 months after the irradiation). Cyst formation was age-dependent but not dose-dependent; tumor formation showed both age-dependence and a dose threshold. Our studies are the first to provide evidence that single sub-lethal TBI can cause long-term ovarian damage like cysts and tumors in mice. The mouse model presented herein may be a useful tool for mechanistic studies on the development of ovarian cysts and tumors, particularly with their shorter life and estrous cycle.

## Materials and Methods

### Mice

The mice used in this study were part of a larger study, which was supported by the grant RFP NIAID-DAIT-NIHAI2008023. According to the original design, a total of 1000 mice were irradiated including males and females. More specifically, 210 C57BL6/J female mice were purchased from The Jackson Laboratory (Bar Harbor, ME) and 200 mice were divided into four age groups. To irradiate the mice, 50 mice at 2 months of age were brought into the radiation room each time, 10 mice in each of 5 groups were exposed to doses of either 0.5, 1, 2, or 4 Gy, with the remaining 10 mice (the 0 Gy group) not exposed to radiation as a control. Six additional mice were exposed to 0.5 or 1 Gy under these same conditions. Therefore, we irradiated a total of 166 mice and had 40 unexposed mice as 0 Gy control. The mice in each dose were subsequently sacrificed at 3.5, 9, 12, or 18 months of age and tissues harvested for assessment. All mice were maintained in a specific pathogen-free facility at the University of Georgia before and after irradiation. The experiment was approved by the Institutional Animal Care and Use Committee of the University of Georgia.

### Irradiator survey and calibration

The radiation exposure device used in this study is a Type 60 Co gamma (Nordion Canada), Model Gammacell 200. To calibrate radiation exposure doses, OSL nanoDot dosimeters (Landauer Inc., Glenwood, IL) were implanted in a mouse cadaver, which were then exposed at the indicated doses of 0.5, 1, 2, and 4 Gy. After irradiation, the dosimeters were sent back to Landauer Inc. for measurement. Based on the results of these measurements, we generated a linear formula to calculate the exposure times for each dose (Y(dose)=2.16t+10.98; S1 Dataset). The standard error of the dose for each irradiation is within 5% [26]. Details of the calibration protocols and evaluation of linear response were previously published [26].

### Measurement of cyst and tumors

Mice in each dose group were sacrificed by CO_2_ inhalation followed by cervical dislocation at 3.5, 9, 12, or 18 months of age and tissues harvested for assessment. The peritoneal and thoracic cavities of the mice were opened, and the ovary, spleen, liver, gut, thymus, lung, and other organs were evaluated for visually apparent anomalies. To measure ovarian cysts and tumors, the ovaries were completely exposed along the uterus on both sides. Two individuals independently assessed and measured cyst formation and tumor lesions. The visible cysts and tumors with a size > 1 mm^3^ and any visible pathological changes in the ovaries and other organs were recorded (S2 Dataset). In this study, the analysis of 3.5-month-old mice at 1.5 months after irradiation was defined as “short-term” (to contrast with “acute”). We defined “long-term” effects as occurring at or after the 9 months age at analysis, more than 6 months after exposure.

### H&E staining

For histological analysis, ovarian samples included unirradiated control (0 Gy), irradiated without visible cysts or tumors, irradiated with visible cysts or tumors were assessed. Tissues were fixed with 4% paraformaldehyde (PFA) overnight, dehydrated in gradient ethanol solutions, and embedded in paraffin. Ten μm sections were cut and stained with hematoxylin and eosin (H&E). Imaging was done on a Keyence BZ-X700 microscope (Osaka, Japan).

### Statistical analysis

One-sided Fisher’s exact tests were used to compare the overall incidence rates between the irradiated and control group. Within the irradiated group, two-sided Fisher’s exact tests were used to assess the dependence of incidence rates across the age and dosage groups respectively. Furthermore, simple logistic regression was used to assess the magnitude and directionality of significant associations with age. These analysis were performed in R using functions from the ‘stats’ package version 3.6.1. (S3 Dataset). The rates of pathological changes in the liver, spleen, lung, thymus, and skin (Table 1) was analyzed using Contingency table analysis, and Fisher’s exact test using Prism™ software (GraphPad Software, San Diego, CA).

**Table 1.** Visible pathological changes after TBI.

|  | Non-irra. | Irradiated groups (all doses) |  |  |  |  |  |
| --- | --- | --- | --- | --- | --- | --- | --- |
| Organ | Ovary <sup>a</sup> | Ovary | Liver | Spleen | Thymus | Lung | Skin |
| No. of total organs examined<sup>b</sup> | 80 | 332 | 166 | 166 | 166 | 166 | 166 |
| No. of organs with cyst | 2 | 49 | 0 | 0 | 0 | 0 | 0 |
| No. of organs with tumor | 0 | 17 | 4 | 3 | 2 | 1 | 1 |
| % Path-change <sup>c</sup> | 2.5% | 19.9% | 2.4% | 1.8% | 1.2% | 0.6% | 0.6% |

Non-irra: Non irradiated group; Path-change: Pathological change
<sup>a</sup> Non-irradiated control mice had no visible changes in any other organs.
<sup>b</sup> As there are two ovaries/mouse, the numbers are double
<sup>c</sup> The overall frequencies of visible pathological changes observed per organ

## Results

### Overall pathological changes in the ovary and other organs after TBI

To evaluate the effects of radiation with aging, two-month-old female mice were total body exposed to a single radiation dose of 0, 0.5, 1, 2, or 4 Gy, and analyzed at 3.5, 9, 12, and 18 months of age. We previously reported that a single TBI could cause long-term deficits in thymus function by reducing lymphoid progenitors in bone marrow [25]. Interestingly, we also found visible pathological changes in the ovary in these mice (Fig. 1). Most ovarian cysts were unilateral, a single round shape, within the bursa, and filled with transparent fluid, with some displaying vascularization (Fig. 1A); a few appeared as large oval shapes formed by multiple cysts (Fig. 1B). Some cysts or masses were bilateral or appeared deep brown or black due to being filled with blood (Fig. 1 C). The ovarian tumors appeared as a large mass of tissue or black masses covered by the bursa (Fig. 1 D). We also observed that some mice had enlarged spleens or tumor lesions in the liver, lung, thymus, and/or skin after TBI. The overall incidence of visible pathological changes after irradiation per ovary was 19.9%; The rates of pathological changes were much lower in the liver, spleen, lung, thymus, and skin (Table 1). Contingency table analysis using Fisher’s exact test showed that pathological changes were significantly different between the ovary and other organs (P<0.001). Our results indicate that the ovary is significantly more sensitive to radiation compared to the other organs examined.

**Fig 1.**
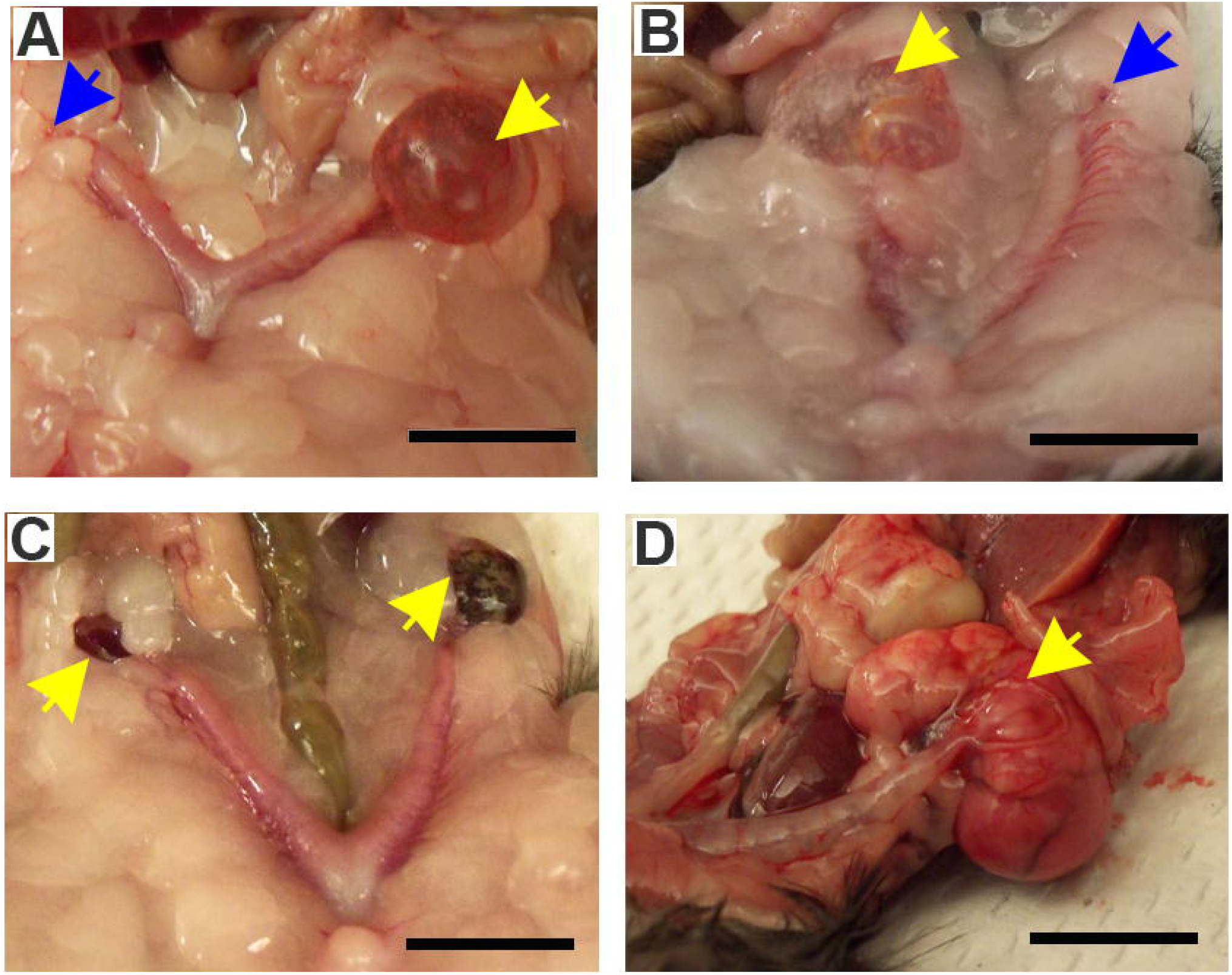
Gross anatomy of ovarian cysts and tumors from BL6 mice after TBI. **(A**). Single round ovarian cysts were filled with transparent or hemorrhagic serous fluid. **(B)**. Oval ovarian cysts filled with transparent fluid and formed by multiple cysts. **(C)**. Ovarian cysts or lumps were filled with hemorrhagic serous fluid. **(D)**. Ovarian tumors were covered with a membrane. Normal ovary (blue arrow); Cysts or tumors (yellow arrow). Size of bars: 10 mm.

### TBI increased cyst formation in the ovary

Next, we measured cyst formation and observed that it is a prominent pathological change in the ovary after TBI. There were no visible cysts observed in either the control or irradiated groups at 3.5 months of age (1.5 months after irradiation), and cysts were observed to be present at 9 months of age (7 months after irradiation) in the irradiated group. The overall rate of cyst formation in irradiated mice is 24.7% compared to 5% in non-irradiated control mice (Table 2, P=0.003 Fisher’s exact test one-sided), indicating that TBI exposed mice were more likely to develop ovarian cysts. Among irradiated mice, we found that the rate of cyst formation was age dependent (Table 2, P<0.001 Fisher’s exact test two-sided). Furthermore, logistic regression showed that age was positively associated with the rate of cyst formation (Odds Ratio=1.15, P<0.001). We also found the rate of cyst formation to be independent of dose (Table 2, P=0.166 Fisher’s exact test two-sided).

**Table 2.** Ovarian cyst formation observed per animal by age and dose after TBI.

| **Age \ Dose** | **Non-irra.** | **Irradiated groups (Gy)** |  |  |  |  |
| --- | --- | --- | --- | --- | --- | --- |
|  | **0** | **0.5** | **1** | **2** | **4** | **% mice with cysts by age** |
| **3.5** | 0<sup>a</sup>/10<sup>b</sup> | 0/10 | 0/10 | 0/10 | 0/10 | 0% (0/40) |
| **9** | 0/10 | 4/10 | 2/10 | 2/10 | 2/10 | 25% (10/40) |
| **12** | 1/10 | 2/10 | 9/12 | 1/10 | 4/10 | 38.1% (16/42) |
| **18 (M)** | 1/10 | 1/12 | 3/12 | 4/10 | 7/10 | 34.1% (15/44) |
| **% mice with cysts by dose** | 5% | 16.7% | 31.8% | 17.5% | 32.5% | **24.7% (41/166)<sup>c</sup>** |

Non-irra: Non irradiated group
<sup>a</sup> Number of mice with cysts
<sup>b</sup> Number of total mice
<sup>c</sup> Proportion of irradiated mice with cyst

### TBI induced tumor lesion in the ovary

Tumor lesions are frequently observed as a long-term consequence after exposure to radiation. Visible tumor lesions (Size > 1 mm^3^) in the ovary were observed to be present at 9 months of age in the irradiated group. The overall rate of tumor lesions in irradiated mice is 10.2% compared to 0% in control mice (Table 3, P=0.024 Fisher’s exact test one-sided), indicating that TBI exposed mice were significantly more likely to develop tumor lesions in the ovary. Among irradiated mice, we found that the rate of tumor lesions was age dependent (Table 3, P<0.001 Fisher’s exact test two-sided). Furthermore, logistic regression showed that age was positively associated with the rate of tumor lesions (Odds Ratio=1.37, P<0.001). Tumors were observed at the doses of 0.5, 1, and 2 Gy respectively, while no tumor lesions were observed in the mice exposed to the 4 Gy dose, suggesting that tumor incidence had a threshold-like effect that was significantly correlated to dose (Table 3, P <0.001 Fisher’s exact test two-sided).

**Table 3.** Ovarian tumor lesions observed per animal by age and dose after TBI.

| Dose<br>Age | Non-irra. | Irradiated groups (Gy) |  |  |  |  |
| --- | --- | --- | --- | --- | --- | --- |
|  | 0 | 0.5 | 1 | 2 | 4 (Gy) | % mice with tumors by age |
| 3.5 | 0<sup>a</sup>/10<sup>b</sup> | 0/10 | 0/10 | 0/10 | 0/10 | 0% (0/40) |
| 9 | 0/10 | 0/10 | 1/10 | 0/10 | 0/10 | 2.5% (1/40) |
| 12 | 0/10 | 0/10 | 2/12 | 1/10 | 0/10 | 7.1% (3/42) |
| 18 (M) | 0/10 | 4/12 | 5/12 | 4/10 | 0/10 | 29.5% (13/44) |
| % mice with tumors by dose | 0% | 9.5% | 18.2% | 12.5% | 0% | 10.2% (17/166)<sup>c</sup> |

Non-irra: Non irradiated group
<sup>a</sup> Number of mice with tumors;
<sup>b</sup> Number of total mice
<sup>c</sup> Proportion of irradiated mice with tumor

### The overall visible pathological changes in the ovary after TBI

To evaluate the radiation-induced effects in the ovary, we combined both cyst formation and tumor lesions to assess pathological changes after TBI in different ages and doses (Table 4). The overall rate of pathological change among irradiated mice is 33.7% compared to 5% in non-irradiated control mice (Table 4, P<0.001 Fisher’s exact test one-sided), indicating that TBI exposed mice were significantly more likely to develop any pathological change. Among irradiated mice, we found that the incidence of pathological change is age dependent (Table 4, P<0.001 Fisher’s exact test two-sided) and the direction of association is positive (Odds Ratio=1.27, P<0.001). We also found that the incidence of pathological change is independent of dose (Table 2, P=0.177 Fisher’s exact test two-sided).

**Table 4.**
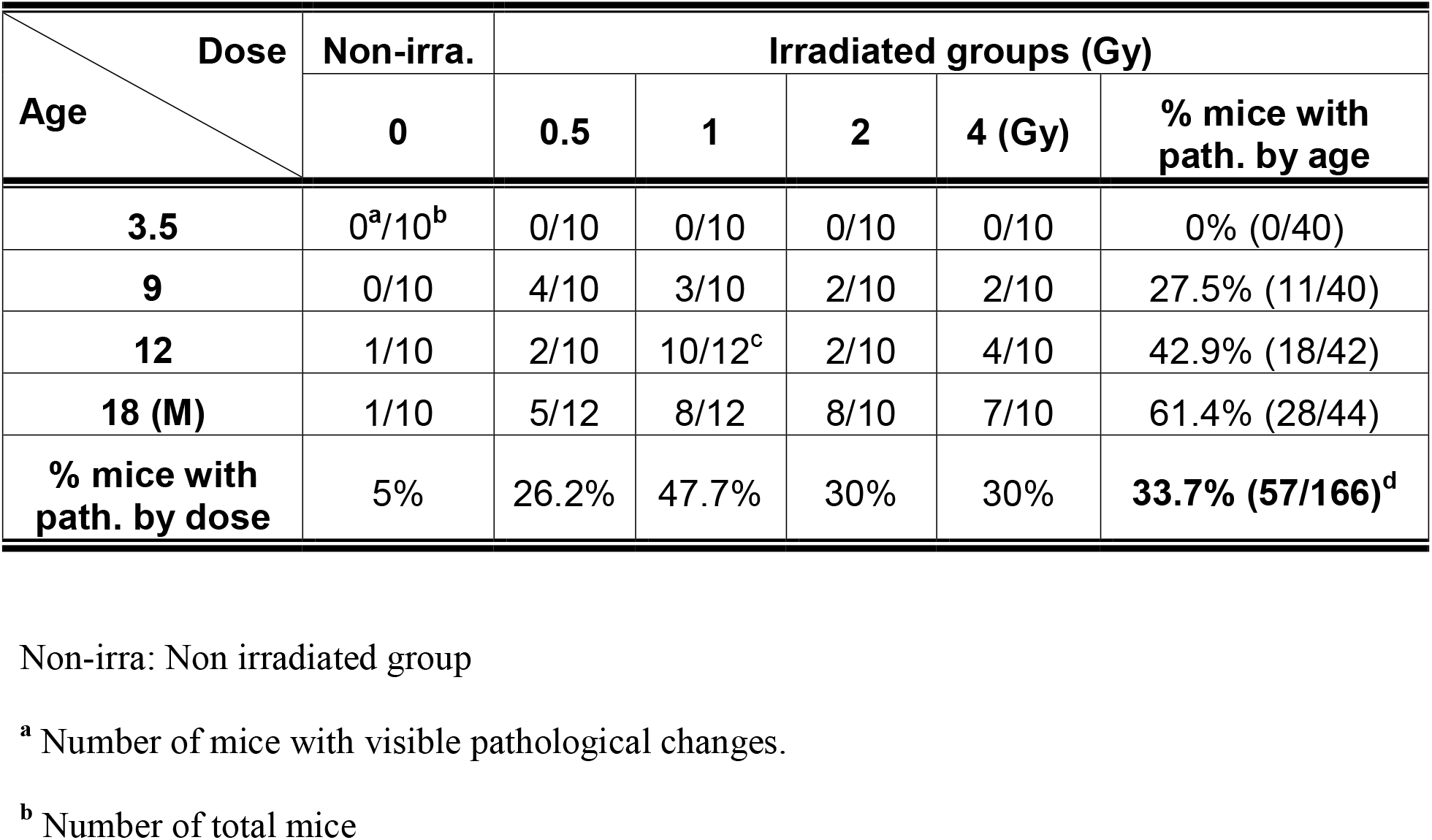
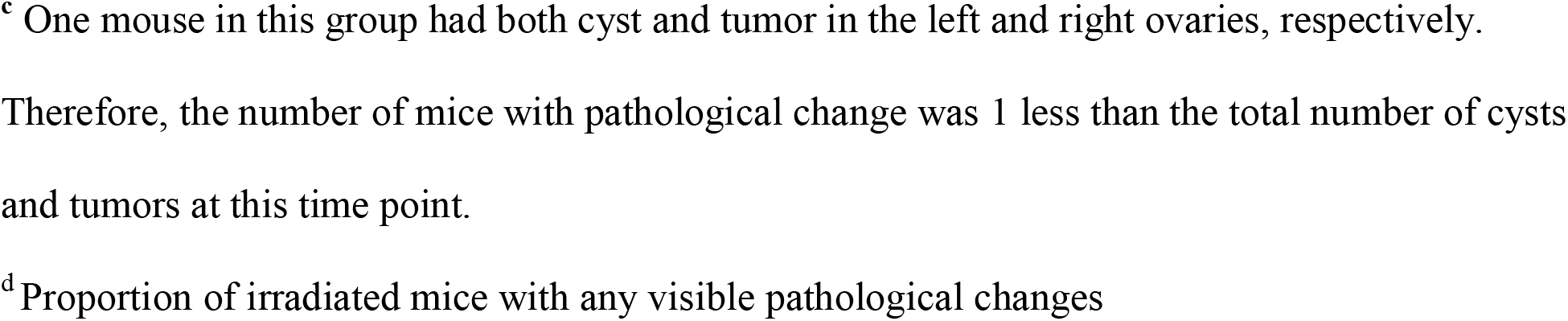
Overall visible pathological changes in the ovary observed per animal by age and dose after TBI.

| Age \ Dose | Non-irra. | Irradiated groups (Gy) |  |  |  |  |
| --- | --- | --- | --- | --- | --- | --- |
|  | 0 | 0.5 | 1 | 2 | 4 (Gy) | % mice with path. by age |
| 3.5 | 0<sup>a</sup>/10<sup>b</sup> | 0/10 | 0/10 | 0/10 | 0/10 | 0% (0/40) |
| 9 | 0/10 | 4/10 | 3/10 | 2/10 | 2/10 | 27.5% (11/40) |
| 12 | 1/10 | 2/10 | 10/12<sup>c</sup> | 2/10 | 4/10 | 42.9% (18/42) |
| 18 (M) | 1/10 | 5/12 | 8/12 | 8/10 | 7/10 | 61.4% (28/44) |
| % mice with path. by dose | 5% | 26.2% | 47.7% | 30% | 30% | **33.7% (57/166)<sup>d</sup>** |

Non-irra: Non irradiated group
<sup>a</sup> Number of mice with visible pathological changes.
<sup>b</sup> Number of total mice
<sup>c</sup> One mouse in this group had both cyst and tumor in the left and right ovaries, respectively.
Therefore, the number of mice with pathological change was 1 less than the total number of cysts and tumors at this time point.
<sup>d</sup> Proportion of irradiated mice with any visible pathological changes

### Both ovarian cyst formations and tumor lesions after TBI originated from granulosa cells

To measure the morphology, structural changes and identify the cells of origin in the cysts and tumors, a subset of ovarian samples with cysts and tumors were collected. The unirradiated ovary and irradiated ovary without visible cysts and tumors were set as controls. These samples were fixed and the paraffin-embedded sections were processed for H&E staining. Under the microscope, the unirradiated, morphological structure of the ovary included clear cortex and medulla areas, whereas there were no follicles in the cortex, which were instead replaced by corpora lutea (CL) (Fig 2 A and B). All irradiated ovaries including non-visible cystic (Fig 2 C and D) and visible cystic ovaries (Fig 2 E-H) had lost the entire structure of cortex, medulla and corpora lutea, but displayed increased cysts in the ovaries. The granulosa cells either surrounded the cyst or were disorderly distributed in the residual ovarian tissues. Most cysts showed follicular origin profiles lined by one or several layers of cuboidal or flattened granulosa cells with a thin wall. Some granulosa cells were luteinized with eosinophilic cytoplasm (Fig 2 C-H blue arrow). Some cysts were located within the ovary, with an epithelial cell cyst profile of a thin wall lined by a layer of flattened epithelium that resembled ovarian surface epithelial cells (Fig 2 C and D, red arrow).

**Fig 2.**
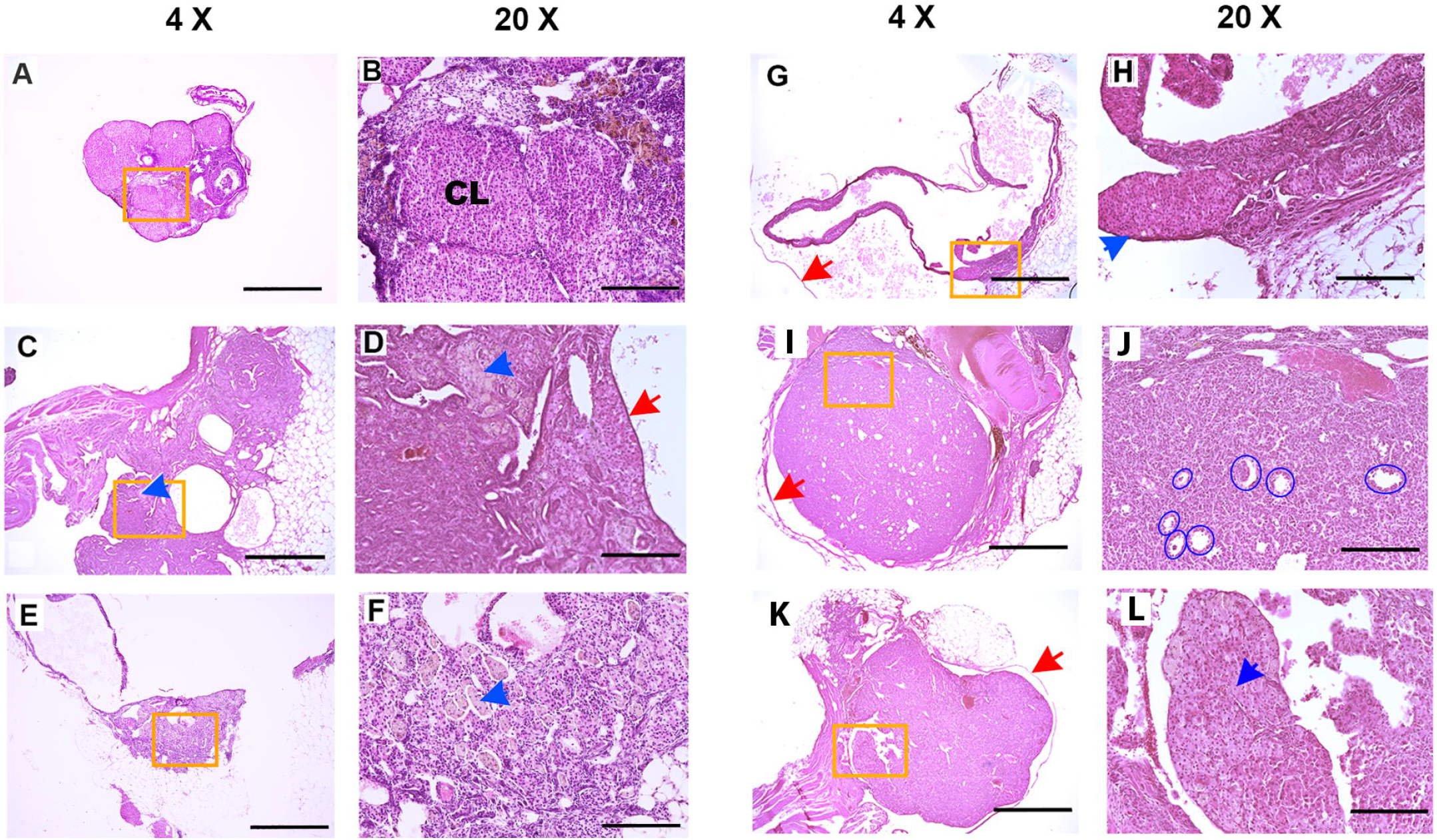
H&E staining profiles of ovarian cysts and tumors. **(A)**. An ovary from an unirradiated 18 month-old mouse as 0 Gy control. **(B)**. Higher magnification image of the indicated area in A, showing cortex with corpora lutea (CL). **(C)**. An ovary without a visible cyst from a mouse exposed to 0.5 Gy, analyzed at 18 months. **(D)**. Higher magnification image of the indicated area in C, showing an ovarian inclusion cyst located in the ovarian tissue lined by a layer of flattened epithelium resembling the ovarian surface (red arrow). **(E)**. A huge follicular cyst located on the ovarian surface with the residual ovary. **(F)**. Higher magnification image of the indicated area in E, showing the disorded structure, cyst and luteinized with eosinophilic cytoplasm granulosa cells (blue arrow). **(G)**. A huge follicular cyst located on the ovarian surface lined by several layers of cuboidal granulosa cells (blue arrow) with a thin wall, surrounded by the ovarian surface membrane (red arrow). **(H)**. Higher magnification image of the indicated area in G, showing several layers of cuboidal granulosa cells. **(I)**. A massive ovarian tumor enriched by granulosa cells. **(J)**. Higher magnification image in the indicated area in I, with Call-Exner bodies (blue circle or oval). **(K)**. Another ovarian tumor, enriched by luteinized granulosa cells. **(L)**. Higher magnification image showing the indicated area of K; luteinized granulosa cells with eosinophilic cytoplasm (blue arrow). All samples were collected from 18 month-old mice. A-B from unirradiated mice. C-K from irradiated mice. Scale bars: A, C, E, G, I, K = 1mm; B, D, F, H, J, L = 200μm

Two massive ovarian tumors were sectioned and stained. Both tumors were composed of cuboidal granulosa cells indicating they were granulosa cell tumors. In the first tumor, some granulosa cells formed structures of the Call-Exner bodies that resemble immature primitive follicles; some granulosa cells lined the cyst and presented in irregular aggregates in the wall (Fig 2 I and J, blue circles). In the second tumor, the granulosa cells were luteinized with moderated amounts of eosinophilic cytoplasm and showed the pale, oval, and angular nuclei in a disorderly arrangement (Fig 2 K and L, blue arrow). Thus, irradiation destroyed the entire morphological structure. In addition, the granulosa cell is a critical cell type involved in the formation of ovarian cysts and tumor lesions over the long-term after TBI.

## Discussion

In this study, we provide novel evidence that ovarian stromal cells such as follicular granulosa cells are highly sensitive to harm from sub-lethal doses of radiation, resulting in irreparable cellular damage that can manifest much later in life. Ovarian cyst formation and tumor lesions are two significant, long-term effects after TBI in mice.

Most ovarian cysts result when mature follicles fail to rupture and release the oocyte during ovulation [17]. Left behind is an organized series of granulosa cells surrounding the oocyte and a fluid-filled antrum. Other types of cysts, including corpus luteum and theca-lutein, result from an exaggerated physiological response, like hormonal overstimulation of the ovary [27]. TBI or pelvis irradiation can destroy all primordial follicles in mice, as well as later-stage follicles and mature follicles. Because maintaining the ovarian environment requires functional interaction between the follicle cells and stromal cells, damage from radiation destroys both ovarian structure and activity during the estrous cycle, including ovulation [28, 29].

Our data suggests that radiation-induced follicle damage at sub-lethal doses entirely destroys the cortex and medulla structures with a disorded granulosa arrangement and leads to follicular cyst formation among aged ovaries after TBI. Our results show that most of these ovarian cysts have follicular profiles and are lined by granulosa cells, indicating that most of these radiation-induced ovarian cysts originated from follicles. Furthermore, cyst formation significantly increased with age, suggesting a long-term impact after TBI includes enhances susceptibility to age-related cyst formation. This is also consistent with estrous cycle changes observed in aging mice, whereby the first phase is characterized by the presence of follicular cysts [30].

Incessant ovulation increases the risk of ovarian inclusion cysts, which are a possible source of epithelial ovarian cancer in women [31]. In this study, we found that although the incidence of ovarian tumors increased after TBI, most of them originated from granulosa cells rather than epithelial cells. We hypothesize that the radiation-induced depletion of ovarian follicles and the subsequent reduction in ovulation may cause this difference. Therefore, these results are consistent with the decreased risk of epithelial ovarian cancer that coincides with ovulation reduction [23, 24]. Ovarian lesions were dramatically more frequent compared to the other organs we evaluated. This observation suggests that the ovary is disproportionally susceptible to damage from radiation.

In contrast with cyst formation, which was age-but not dose-dependent, tumor formation was both and age- and dose-dependent. The dose-dependent effect was mainly driven by in the 4 Gy group, which showed no tumors, and thus was a threshold effect, rather than a linear or exponential dose dependency as has often been reported for other tissues/organs [32, 33]. Since higher radiation doses may directly cause cell death [2, 3], the 4 Gy dose may be sufficient to directly kill sensitive ovarian follicle cells, including oocytes and granulosa cells [7].

While the ovarian tumors incidence was very low overall, we observed a much higher incidence of cysts at 9 and 12 months of age. Numerous studies support that invaginated or trapped inclusion cysts are a possible source of epithelial ovarian cancer and might increase the risk of the disease [34, 35]. In this study, some of the H&E sections showed inclusion cysts, but we did not see tumors of ovarian epithelial origin. Both tumors we analyzed originated from ovarian granulosa cells; all profiles indicate that they were ovarian granulosa cell tumors [36-38]. Granulosa cell tumors are a rare type of slow-growing ovarian cancer, a part of the sex cord-gonadal stromal tumor that accounts for 2% to 5% of all malignant ovarian cancers [39, 40]. Therefore, our data suggest that sublethal radiation damage is less likely to induce epithelial ovarian cancer, but instead can induce granulosa cell tumors.

To summarize, in the present study, we provide new evidence that the ovary is disproportionately sensitive to radiation, even at relatively low doses. Ovarian cyst formation and tumor lesions both occur as long-term effects after TBI in mice. Granulosa cells are the primary target cells that contribute to these changes. Because most cysts originate from follicles, this mouse model may provide a way to analyze the underlying mechanisms of ovarian cyst formation caused by radiation-induced oocytes/follicular cell damage, hormone imbalance, and ovulation cycle disorders. Also, because we have no human clinical trial data to evaluate the genetic effects of human exposure to ionizing radiation, extrapolating human genetic risk is based on mouse data. Our studies may provide a reference for the clinical protection of radiation therapy in females. However, we should note that because the radiation sensitivity of follicle cells varies widely depending on stages and species [8, 15], extrapolation of the ovarian data from mouse to humans requires caution.

## Acknowledgments

We thank Dr. Scott Noakes (Center for applied isotope studies, University of Georgia) for setting up the irradiation of mice. And thank Dr. Thomas Seed (Tech micro services Co, Bethesda, MD) for the irradiator calibration.

## Authorship and Contributions

S.X. designed the study, performed all experiments, analyzed data, and wrote the manuscript. Y.X. performed statistical analysis on the data. X.X. analyzed H&E staining images. W.Z. helped to perform all experiments and took care of mice. M.M.M. participated in discussions about the data and wrote the manuscript. N.R.M. designed the study and discussed the results and wrote the manuscript. All authors read and edited the manuscript before submission.

## Supporting information

**S1 Dataset. Calibration of radiation dose rate**.

**S2 Dataset. Sample record of pathological changes in ovaries and other organs**.

**S3 Dataset. Report of statistical analysis result**.

